# Host adaptive radiation is associated with rapid virus diversification and cross-species transmission in African cichlid fishes

**DOI:** 10.1101/2023.06.28.546811

**Authors:** Vincenzo A. Costa, Fabrizia Ronco, Jonathon C.O. Mifsud, Erin Harvey, Walter Salzburger, Edward C. Holmes

**Author notes:** Corresponding author: Edward C. Holmes, Sydney Institute for Infectious Diseases, School of Medical Sciences, The University of Sydney, Sydney, NSW 2006, Australia.

## Abstract

Adaptive radiations are generated through a complex interplay of biotic and abiotic factors. Although adaptive radiations have been widely studied in the context of animal and plant evolution, little is known about how they impact the evolution of the viruses that infect these hosts, which in turn may provide insights into the drivers of disease emergence. We examined how the rapid adaptive radiation of the African cichlid fishes of Lake Tanganyika over the last 10 million years has shaped the diversity and evolution of the viruses they carry. Through metatranscriptomic analysis we identified 121 vertebrate-associated viruses among various tissue types that fell into 13 RNA and 4 DNA virus groups. Host-switching was commonplace, particularly within the *Astroviridae*, *Metahepadnavirus*, *Nackednavirus*, *Picornaviridae*, and *Hepacivirus* groups, occurring more frequently than in other fish communities. A time-calibrated phylogeny revealed that hepacivirus evolution was not constant throughout the cichlid radiation, but accelerated 2-3 million years ago, coinciding with a period of rapid cichlid diversification and niche packing in Lake Tanganyika, thereby providing more closely related hosts for viral infection. These data show that African cichlids contain a complex interacting pool of virus diversity, likely reflecting their close genetic relationships that lowers the barriers to cross-species virus transmission.

---

Since the origin of vertebrates some 540 million years ago (Ma), the evolution of the viruses that infect these animals has been shaped by a combination of host-virus co-divergence over many millions of years coupled with cross-species virus transmission (i.e., host-switching) events that occasionally result in disease outbreaks^1–7^. Although viruses regularly emerge in new hosts, there is increasing evidence that viruses are natural and abundant components of ecosystems and may be transmitted among species with no apparent disease^8–10^. Teleost fishes, the largest clade of living vertebrates that display extensive ecological diversity, are among those groups that naturally exhibit highly diverse viromes, making them an informative model system for determining the host and virological factors that impact cross-species virus transmission^10, 11^. Indeed, relatives of coronaviruses, hepatitis C virus, influenza viruses, filoviruses, and paramyxoviruses have all been identified in teleosts^2, 10–15^.

Adaptive radiations—the rapid formation of a species complex through adaptations to new or existing ecological niches—have generated substantial biodiversity in teleost fishes, with iconic examples including the three-spined sticklebacks, Antarctic notothenioids, and most notably the cichlid fishes inhabiting the East African Great Lakes, Tanganyika, Malawi, and Victoria^16–18^. The African cichlid radiations rank among the most rapidly evolving species flocks in vertebrates, comprising nearly 2000 species, all of which have evolved in less than 10 million years^19–21^. The cichlid radiation in Lake Tanganyika is the oldest of these flocks, with approximately 240 endemic species arranged into 12 phylogenetic subclades (i.e., “tribes”)^19, 20^. Despite their phenotypic diversity, African cichlids exhibit lower levels of genetic diversity than most non-radiating vertebrate populations, and many alleles are shared among species from different lakes as well as between those with strikingly different morphologies, body sizes, and ecologies^22^. From a virological perspective, the limited genetic diversity among Tanganyikan cichlids should also mean that these animals will exhibit broadly similar cell receptors, in turn lowering the adaptive barriers for viruses to jump species boundaries. Conversely, it is possible that viruses might co-diverge with the rapidly evolving cichlids into different ecological niches. To date, these ideas have not been tested.

Determining the viromes of African cichlids can also provide insights into how key host transitions have shaped the macro- and microevolution of viruses. The cichlid radiation began in Lake Tanganyika approximately 9-10 Ma^20, 23^ and the divergence times of these host species can be used to estimate the rate of virus diversification and determine whether and how it has been shaped by major events in host evolution^24^. For instance, as host populations become denser and more diverse, it is expected that the rate of virus evolution, including the rate of virus speciation, will similarly increase. This question has yet to be addressed for viruses.

Here, we examine the relationship between host adaptive radiation and virus evolution, using the exceptionally diverse cichlid fauna of Lake Tanganyika as a model system. In particular, we sought to determine whether viruses have co-diverged with the radiating cichlids or whether there has been more frequent cross-species transmission within a restricted geographical space. In addition, we aimed to reveal the rate of virus diversification in cichlids and assess whether it was constant throughout cichlid evolutionary history, or whether it peaked during periods of rapid host diversification.

## Results

### The Tanganyikan cichlid virome

We screened 23.7 billion RNA sequencing reads (median 62,909 assembled contigs per library) from 74 Lake Tanganyikan cichlid species (n = 6 replicates per species) for vertebrate-associated viral transcripts. The sequencing libraries screened were generated from a previous study of host gene expression dynamics^25^ and are available on the NCBI Sequence Read Archive (SRA) under BioProject PRJNA552202. The taxonomic sample covers all 12 tribes of the cichlid adaptive radiation in Lake Tanganyika: Bathybatini, Benthochromini, Boulengerochromini, Cyphotilapiini, Cyprichromini, Ectodini, Eretmodini, Lamprologini, Limnochromini, Perissodini, Trematocarini, and Tropheini/Haplochromini.

We identified 121 likely vertebrate-associated viruses in 52 cichlid species that fell within the following taxonomic groups (virus genus or family): *Hepacivirus* (33 viruses), *Flavivirus* (7), *Astroviridae* (11), *Matonaviridae* (5), *Caliciviridae* (1), *Picornaviridae* (5), *Coronaviridae* (3), *Hepeviridae* (2), *Nanghoshaviridae* (2), *Piscichuvirus* (4), *Arenaviridae* (1), *Paramyxoviridae* (2), *Birnaviridae* (2), *Ichthamaparvovirus* (2), *Adomaviridae* (7), *Metahepadnavirus* (19), and *Nackednavirus* (15) (Figure 1b; Supplementary Table 1). Among the tribes, we found a positive relationship between the number of cichlid species examined and the number of viruses detected (*R* ^2^*=* 0.74, *P* < 0.001), with the Lamprologini harbouring 40% of the individual viruses identified, followed by the Ectodini (30.4%), Tropheini/Haplochromini (9.6%), and Cyprichromini (8%), reflecting the species richness of these tribes in Lake Tanganyika^19^.

**Figure 1.**
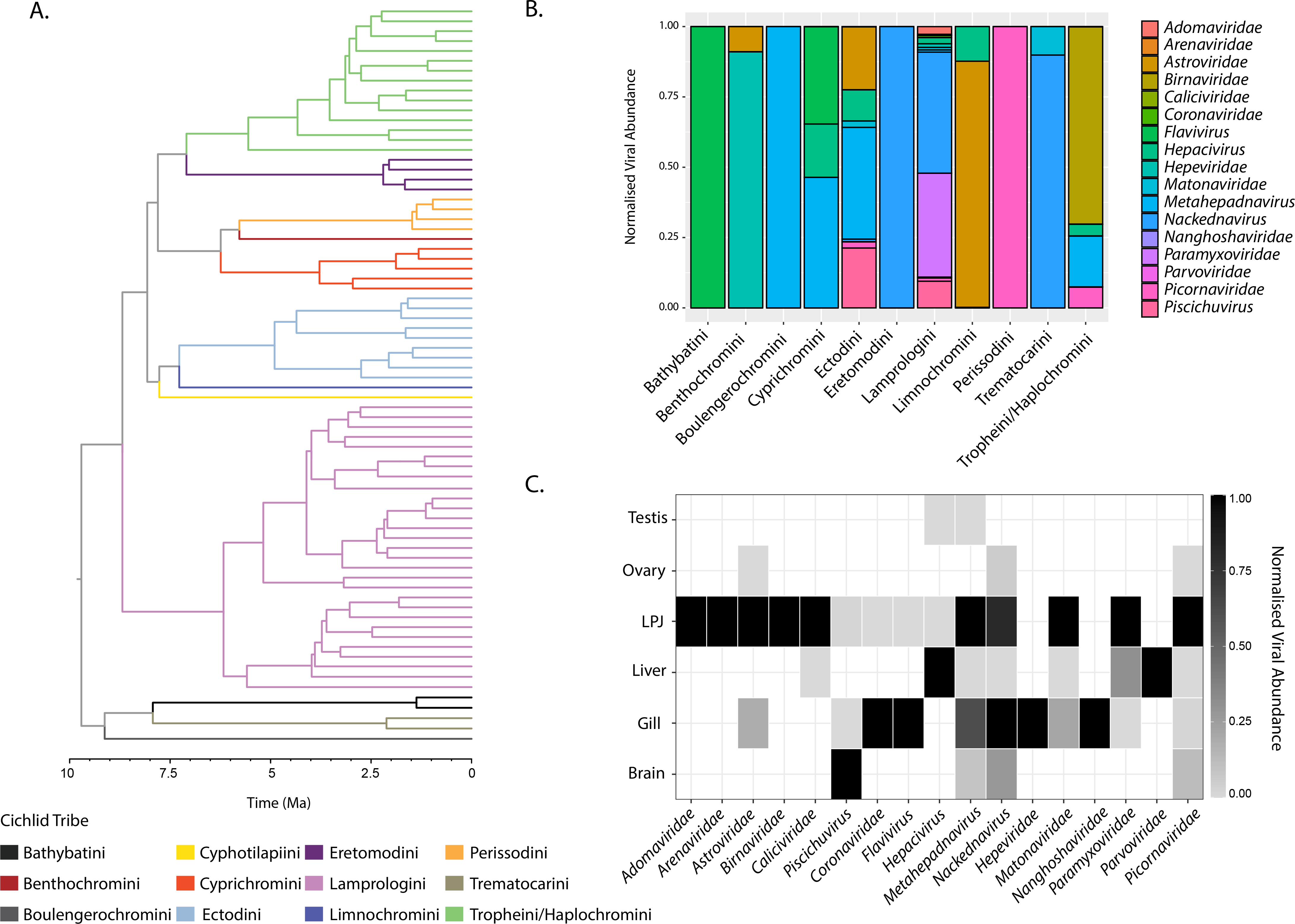
Abundance and diversity of cichlid viromes. **a,** Time-calibrated phylogeny of the cichlid (host) adaptive radiation in Lake Tanganyika. The phylogeny was obtained from the literature^20^ and pruned to the species examined in this study (see Supplementary Figure 1 for host species labels). Tree branches are coloured according to cichlid tribe. **b,** Normalised viral abundance of cichlid tribes. **c,** Normalised viral abundance for each organ.

To explore the potential tissue tropisms of the cichlid viruses identified here, we calculated the transcript abundances of viruses identified in the brain, gill, liver, lower pharyngeal jaw (LPJ; an important component of the cichlid’s feeding apparatus), ovary, and testis. This revealed that 94% of the viral groups were present in the gills and the LPJ, implying that these organs, which are in direct contact with the aquatic environment, are a common transmission route among cichlid viruses (Figure 1c). While the cichlids examined displayed no overt signs of disease, we detected 41 viruses in multiple organs, suggestive of systemic infection (Supplementary Table 1). Notably, the piscichuviruses were predominantly found in brain tissue, while the hepaciviruses showed a higher prevalence in the liver.

### Genome organisation of cichlid viruses

The organisation of cichlid virus genomes were highly conserved, resembling those of other fish viruses and, in many instances, those of their mammalian counterparts (Figure 2). For example, all positive-sense single-stranded RNA viruses had the same number of open reading frames (ORFs) and conserved domains as related mammalian viruses, with the picornaviruses containing a polyprotein with the rhv, helicase, proteinase, and RdRp domains, astroviruses comprising three ORFs, and the hepeviruses and matonaviruses containing a non-structural polyprotein and structural capsid protein, with the latter similar to that of rubella virus (Figure 2a)^26^.

**Figure 2.**
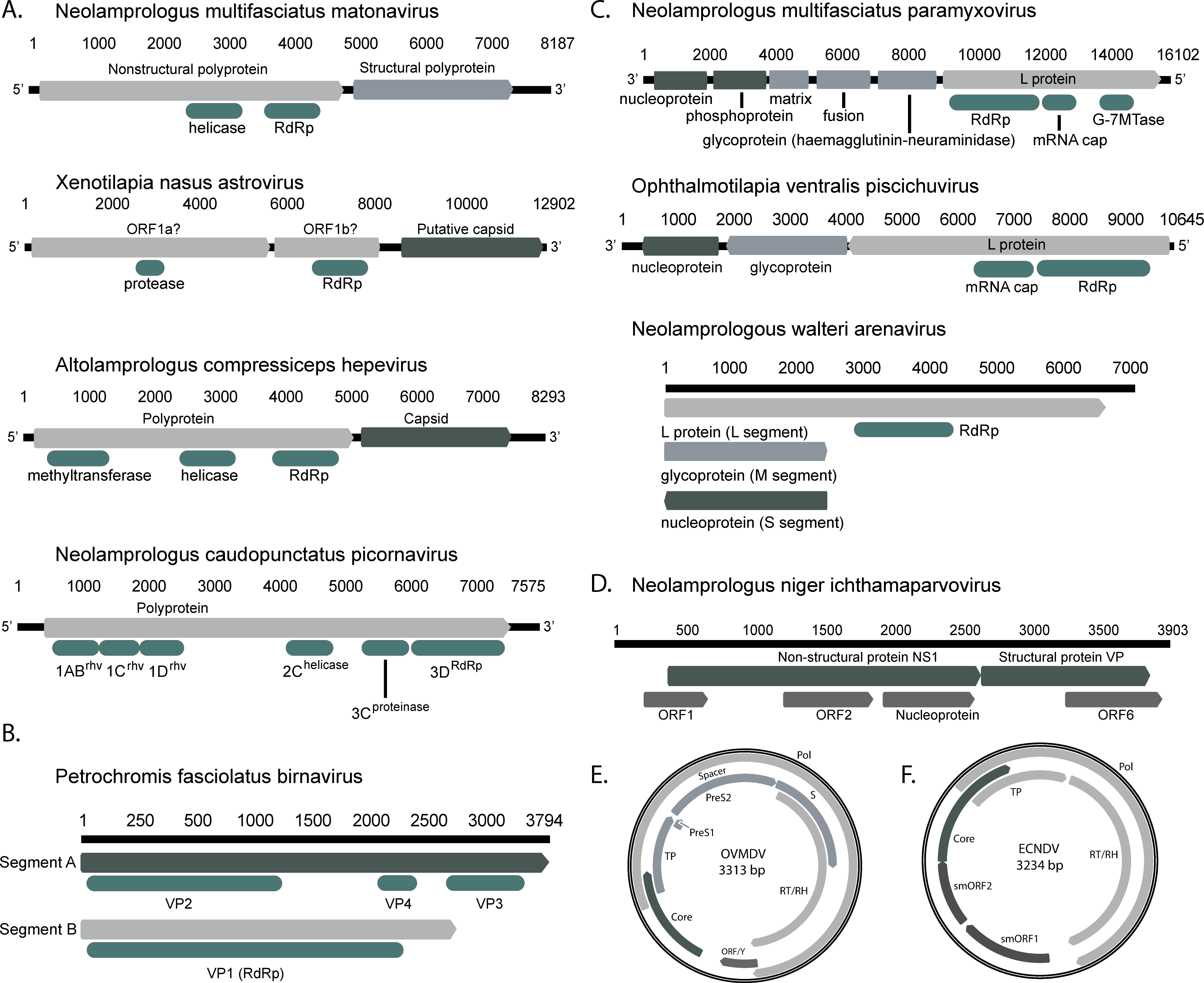
**a,** Genome organisation of positive-sense single-stranded RNA viruses. All viruses are in the 5’-3’ orientation. **b,** Organisation of *P. fasicolatus* birnavirus (double-stranded RNA), showing segments A (structural) and B (RdRp) with VP domains. **c,** Organisation of negative-sense single-stranded RNA viruses. The paramyxovirus and piscichuvirus are shown in the 3’-5’ direction, with the direction of the ambisense arenavirus segments indicated by arrows on bar tips. **d,** ORFs of the single-stranded DNA parvovirus genome, encoding NS1 and VP structural proteins. **e,** Genome of cichlid metahepadnaviruses. OVMDV = *Ophthalmotilapia ventralis* metahepadnavirus. **f,** Genome of cichlid nackednaviruses, with the notable lack of the S (envelope) proteins. ECNDV = *Eretmodus cyanostictus* nackednavirus.

Similarly, the genome of Neolamprologus multifasciatus paramyxovirus had the same order and number of proteins as the orthoparamyxoviruses^27^ (Figure 2c). The N protein encoded an RNA-binding nucleocapsid domain and the three membrane-associated proteins contained a viral matrix protein, fusion glycoprotein domain, and conserved receptor binding protein encoding a haemagglutinin-neuraminidase (HN) domain with a predicted β-propeller fold consisting of six anti-parallel β-sheets (i.e., “propeller blades”)^28^. The second propeller blade had a starting sequence of “HRKSCA”, resembling that of the “NRKSCS” hexapeptide motif (i.e., sialic-acid binding site) found in the respiroviruses, jeilongviruses, ferlaviruses, orthorubulaviruses, and avulaviruses that use sialic acids for cell entry, suggesting that this mode of entry originated in the aquatic environment^27, 28^.

Among DNA viruses, all hepadnaviruses had circular and intact genomes, with the expected length of approximately 3.2 kb, strongly implying that these are exogenous viruses (Figure 2e). The metahepadnaviruses exhibited typical their genome organisation with the viral P protein containing the terminal protein (TP), reverse transcriptase and RNase H domains (RT/RH) as well as the viral core (C) protein and envelope glycoproteins (PreS/S)^29, 30^. All nackednaviruses contained viral C and P proteins, with the notable absence of an envelope^30^ (Figure 2f).

### Ecology and evolutionary history of Tanganyikan cichlid viruses

To determine whether host-switching has shaped the evolutionary history of cichlid viruses, or if it reflects the co-divergence of viruses and cichlids into different ecological niches during the Tanganyikan cichlid radiation, we performed a phylogenetic reconciliation analysis. Accordingly, virus and host phylogenetic trees were mapped against each other, from which we calculated the proportion of the following biological events: co-divergence, duplication, extinction, and host-switching^1, 31^. We focused on the *Hepacivirus, Astroviridae, Matonaviridae, Adomaviridae, Picornaviridae, Piscichuvirus, Metahepadnavirus*, and *Nackednavirus* groups as these contained viruses from multiple cichlid species. This analysis revealed that host-switching was the most common event among virus-host associations (*χ*^2^ = 141.52, *P* < 0.00001), representing 60% of all events, followed by co-speciation (23%), duplication (15%), and extinction (4%). Cross-species virus transmission was particularly common in the astroviruses (72% host-switching events), hepaciviruses (70%), nackednaviruses (65%), metahepadnaviruses (63%), and picornaviruses (61%). Hence, cross-species transmission has played a major role in shaping cichlid virus evolution, occurring in both RNA and DNA viruses (Figure 3).

**Figure 3.**
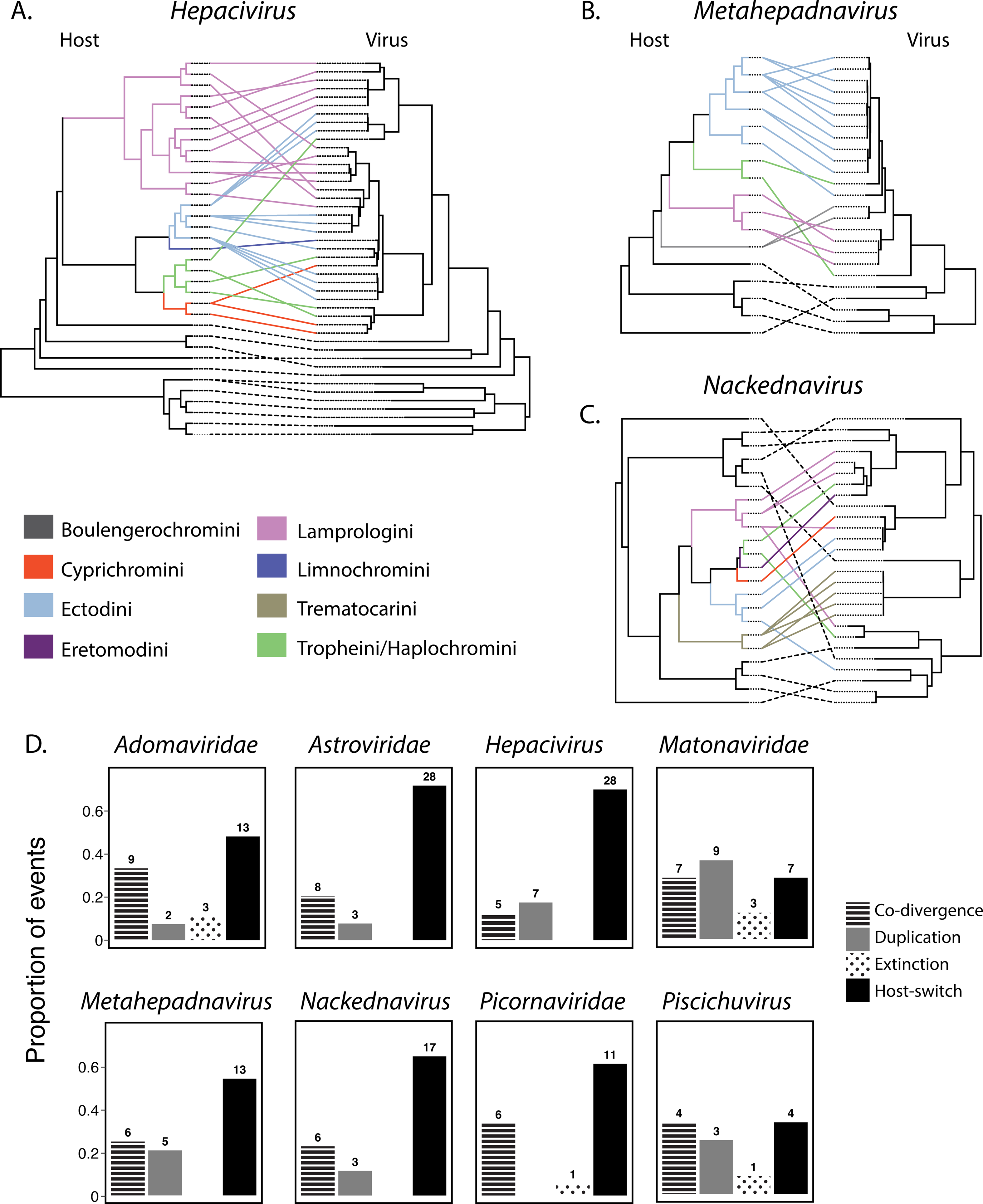
**a,b,c,** Co-phylogenetic analysis of the genera *Hepacivirus*, *Metahepadnavirus*, and *Nackednavirus*, with the host phylogeny on the left and virus phylogeny on the right. Lines connecting taxa are coloured according to cichlid tribe. Dashed black lines represent background sequences of non-cichlid fish viruses. **d,** Phylogenetic reconciliation analysis of cichlid viral groups using eMPRess. Plots show the proportion of events.

To further investigate the impact of host relatedness and ecology on virus evolution, particularly whether cichlid viruses are associated with specific host phylogenetic lineages or ecological niches, we evaluated the phylogenetic and ecological diversity of the infected and uninfected host species based on the host phylogeny and available carbon (δ^13^C) and nitrogen (δ^15^N) stable-isotope measurements. Stable-isotope composition is a reliable measure of benthic-pelagic positioning (δ^13^C) and trophic level (δ^15^N) among fish communities and therefore a strong indicator of host ecological diversity^20^. For instance, fishes with high δ^13^C and low δ^15^N occur mainly in the benthic and/or littoral zone and are lower in the food chain (e.g., littoral herbivores), whereas those with low δ^13^C and high δ^15^N occur in the pelagic or deep zone and have a higher trophic level (e.g., pelagic predators)^20^. Overall, we found a broad distribution of viruses among ecological niches and across the host phylogeny, with no clear ecological or phylogenetic partitioning for all or individual viruses (Supplementary Figure 2a-b). For example, the hepaciviruses, hepadnaviruses, picornaviruses, and astroviruses were found in both pelagic and benthic zones with no structuring according to host ecology. We further compared the ecological and phylogenetic diversity (measured as mean pairwise distances) of infected and uninfected species in our dataset to 1000 subsets of species that we randomly sampled across the cichlid radiation using a one-tailed test, which under the null hypothesis assumes that virus infectivity and transmission is random. This analysis showed that both groups represented random subsets of the radiating cichlids that are not restricted to certain areas in the host tree (infected, *P =* 0.097; uninfected, *P* = 0.796), nor to certain ecological niches (infected, *P =* 0.063; uninfected, *P* = 0.275) (Supplementary Figure 2c). However, we found slightly lower phylogenetic diversity in the infected hepacivirus group compared to random subsets of equal sample size (*P* = 0.049), which may be driven by the presence (and transmission) of these viruses among multiple lamprologine cichlid species (see below). Collectively, these data strongly suggest that cichlid viruses are generalists in Lake Tanganyika and may be readily transmitted among species, irrespective of their ecological niche and, in most cases also independent of the host’s phylogenetic position.

### Close phylogenetic relationships of cichlid RNA viruses

We identified 77 RNA viruses likely associated with vertebrate hosts. Of these, 28% were associated with recent cross-species transmission as the viruses in question were very closely related (>99.5% amino acid similarity). A case in point were the hepaciviruses which fell into three clades (Figure 4). While there was a relatively high level of genetic divergence between these clades (e.g., 60-66% amino acid similarity between clades I and II), there were multiple instances of cross-species virus transmission within individual clades. Within clade I, we identified four viruses that had 99-100% pairwise identity in the NS5 gene in *Julidochromis ornatus*, *Neolamprologus savoryi*, *Neolamprologus furcifer*, and *Chalinochromis brichardi*. We also detected African cichlid picornavirus in five species from the tribes Lamprologini (2 individuals), Perissodini (1), Ectodini (1), and Tropheini (1), as well as African cichlid piscichuvirus in the brain tissue of three species (Figure 4). Similarly, all three piscichuviruses shared 99.2% L protein (RdRp), 98.4% M protein (glycoprotein), and 98.9% S protein (nucleoprotein) similarity. We also identified African cichlid hepevirus in *Altolamprologus compressiceps* and *Benthochromis horii*, and African cichlid flavivirus in two Cyprichromini and *Bathybates fasicatus* (Bathybatini) (Figure 4). Among other RNA viruses, 63% fell within the same genus or species as another cichlid virus, such that they represent a mixture of common ancestry and past host-switching throughout evolutionary history (Figure 4). For example, we identified a novel group of five rubella-like viruses (*Matonaviridae*) that had ∼80% similarity and two birnaviruses that exhibited 93% similarity. While the remaining RNA viruses were divergent (or identified in a single host species), they exhibited phylogenetic relationships to other fish viruses (e.g., *Paramyxoviridae*, *Arenaviridae*) (Supplementary Figure 3).

**Figure 4.**
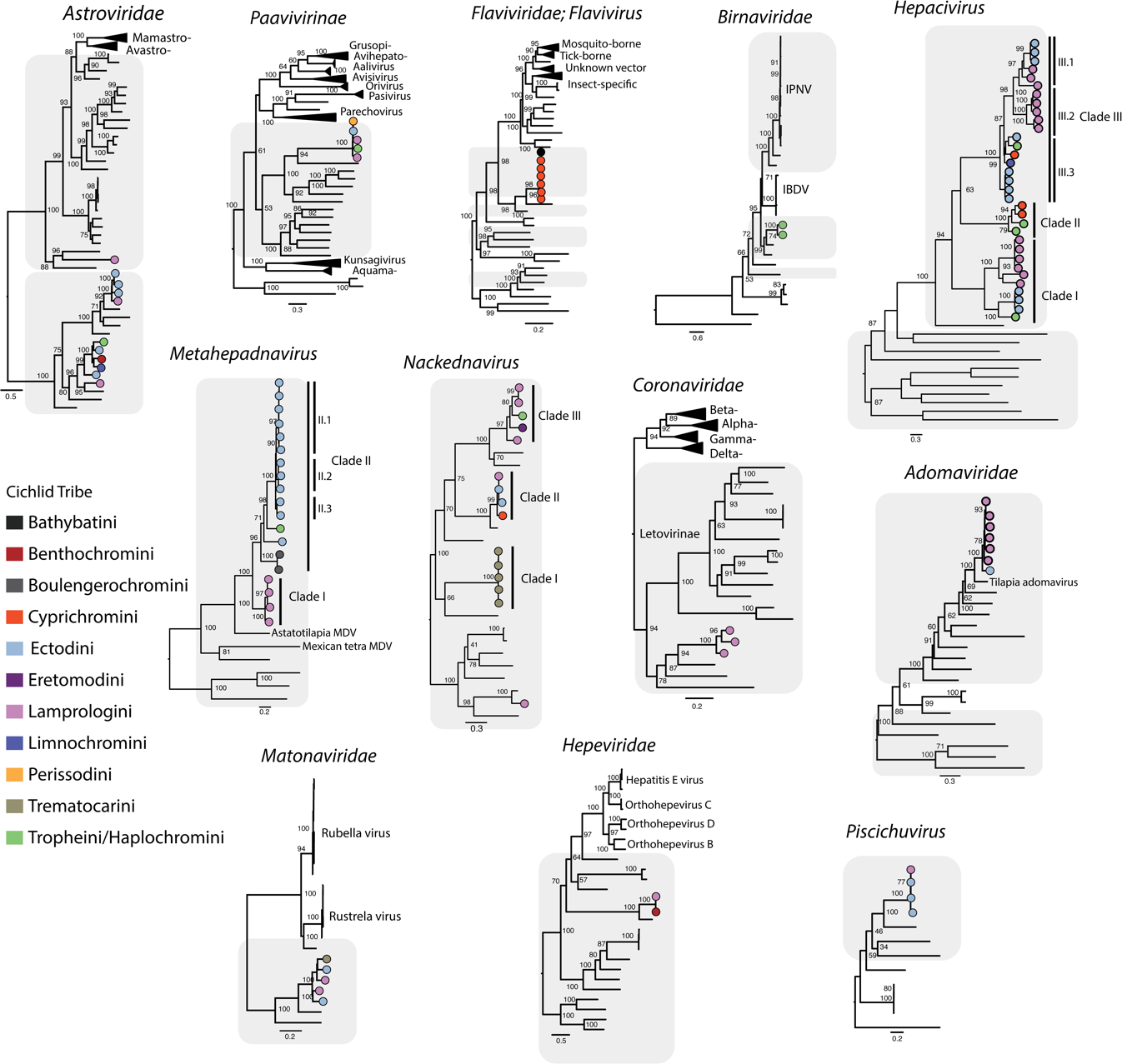
Close evolutionary relationships among cichlid viruses. Phylogenetic analysis reveals close evolutionary relationships among cichlid viruses – in most cases forming monophyletic groups – reflecting recent common ancestry and cross-species virus transmission events. Phylogenies were estimated using amino acid sequences of the RdRp gene for RNA viruses, P protein for the hepadnaviruses, and LO7 for the adomaviruses. Trees were midpoint rooted for clarity only. Circles on branch tips represent individual viruses identified in this study and coloured according to tribe. *Birnaviridae* IPNV = Infectious pancreatic necrosis virus; IBDV = Infectious bursal disease virus. Shaded branches show fish viruses. The scale bar represents amino acid substitutions per site.

### Host-switching and phylogenetic relationships of cichlid DNA viruses

As well as RNA viruses, our analysis identified 43 vertebrate DNA viruses, the majority of which fell within the *Hepadnaviridae*: 19 metahepadnaviruses and 15 nackednaviruses (Figure 4). With the exception of *Lamprologus ocellatus* nackednavirus, all other cichlid hepadnaviruses clustered together with other cichlid viruses in our phylogenetic trees, reflecting cross-species transmission or recent common ancestry (Figure 4). This was particularly apparent in the case of metahepadnaviruses, with examples of cross-species virus transmission occurring within clade II (subclades II.1, II.2, II.3) and clade I (between *Telmatochromis temporalis* and *Lamprologus lemairii*). This pattern was also observed within clades I and II of the nackednaviruses (Figure 4).

We also identified a novel group of adomaviruses that had ∼70% amino acid similarity with tilapia adomavirus^32^, identified in *Oreochromis niloticus*, that shares a common ancestor with Tanganyikan cichlids about 16 Ma^33^ (Figure 4). The high levels of sequence similarity within the adomaviruses (93%) strongly suggests that these viruses were associated with cross-species transmission within Lake Tanganyika (Figure 4).

### Temporal dynamics of *Hepacivirus* evolution in Lake Tanganyika

To infer the temporal dynamics of cichlid virus evolution in Lake Tanganyika we performed a molecular clock analysis using the hepaciviruses as a model system. We based our analysis on this group as they contained the largest number of viruses (n = 33), are monophyletic, and follow a broad pattern of virus-host co-divergence. For example, the genus *Hepacivirus* includes a monophyletic fish group^13^ that broadly reflects the phylogenetic history of its hosts; the ray-finned fish viruses (Actinopterygii) share a common ancestor with a lungfish hepacivirus (Sarcopterygii) and more divergent viruses are associated with cartilaginous fishes (Chondrichthyes).

To determine the rate of hepacivirus diversification in Lake Tanganyika, we estimated the divergence times of all currently described ray-finned fish hepaciviruses using prior age estimates of the formation of Lake Tanganyika, dated at 9-12 Ma^23^, which also reflects the age of the cichlid radiation that was recently estimated at 9.7 Ma^19, 20^. Our calibration was therefore placed at the root node of the cichlid hepacivirus clade with an age interval of 9-12 Ma, following its earlier divergence from *Chaetodon aureofasciatus* hepacivirus^10^. This analysis revealed that the Percomorpha hepacivirus clade had a mean divergence time of approximately 5 million years from other ray-finned fish hepaciviruses (95% higher posterior density [95% HPD]: 3.5-6.9 million years) (Figure 5), while the most recent common ancestor (MRCA) of the cichlid hepaciviruses had a mean age of ∼1.6 million years ago (95% HPD: 0.7-2.7 million years).

**Figure 5.**
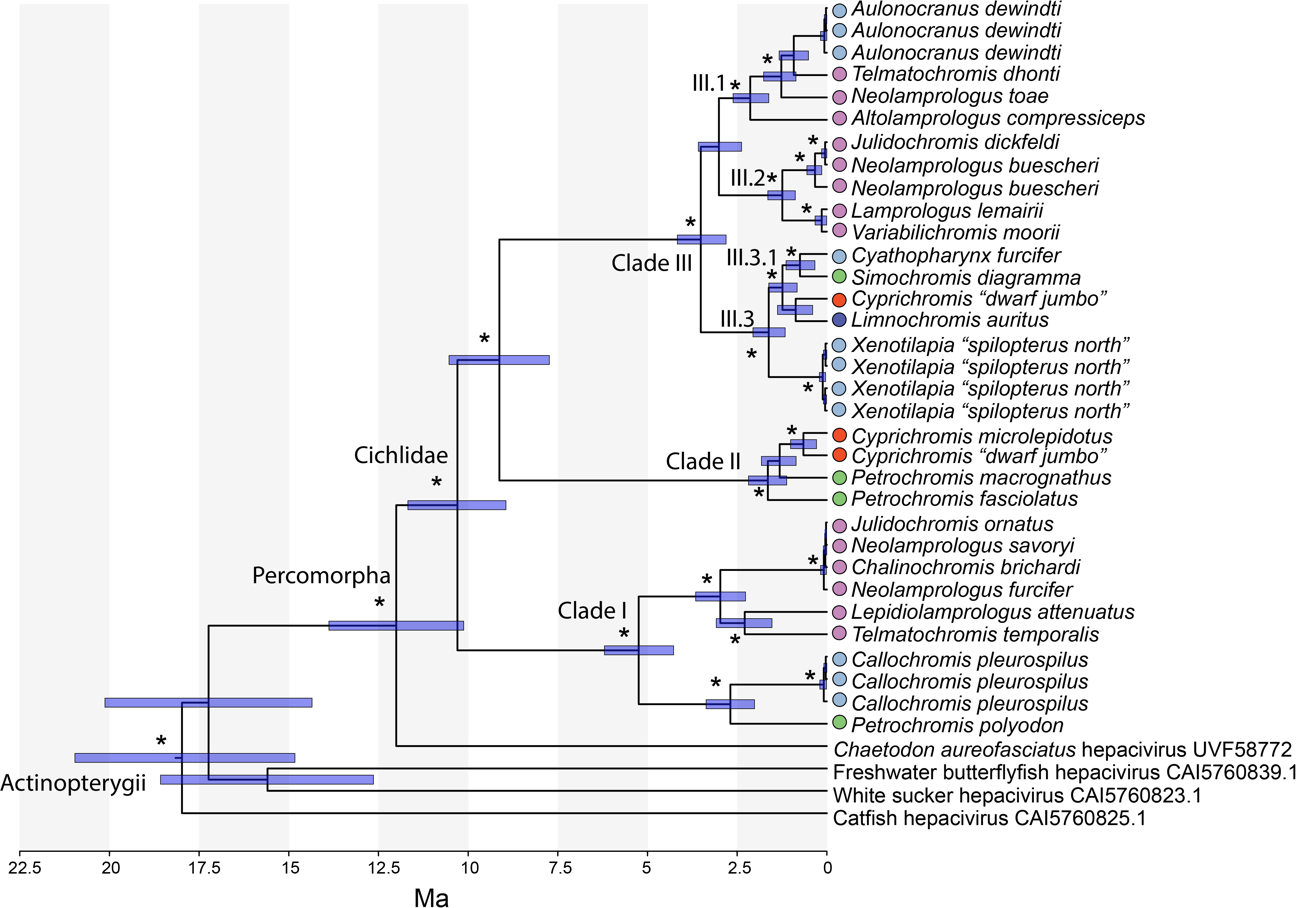
Time-calibrated maximum clade credibility tree of ray-finned fish (Actinopterygii) hepaciviruses. The time axis represents millions of years ago (Ma). Node bars represent the 95% HPD interval for age. Stars denote a posterior probability of over 0.85. Tip labels are coloured according to cichlid tribe.

To test whether the rate of cichlid hepacivirus diversification changed over time we computed the gamma statistic (γ), the null model of which assumes that rates of clade diversification have remained constant over evolutionary history^34^. Accordingly, our time-calibrated phylogeny revealed that the rate of hepacivirus diversification was not constant throughout the cichlid radiation (γ = 3.864, *P =* 0.001), but seemingly began to accelerate around 2-3 Ma, with multiple viral species forming during the last 1-2 Ma (Figure 5). Overall, this coincides with the most recent of three pulses of cichlid evolution in Lake Tanganyika, a period of rapid host diversification^19^.

## Discussion

Evolutionary radiations are responsible for much of the earth’s biodiversity and often involve complex interactions between both biotic and abiotic factors^35^. The interplay between these factors is also central to the generation of virus diversity: viruses routinely adapt to genetic and immunological barriers when they invade new hosts, and there must be adequate exposure between host species for intra- and inter-host transmission^36^. By analysing the viromes of 74 closely related cichlid species endemic to Lake Tanganyika, we show for the first time that host adaptive radiation is associated with rapid virus diversification and cross-species transmission. From a total of 121 vertebrate-associated DNA and RNA viruses identified, 94% were characterised by both recent common ancestry and host-switching.

The evolutionary patterns observed here support the idea that viruses more readily jump between closely related host species, perhaps because of an underlying similarity in their cell receptors which in turn lowers the adaptive barriers to viruses when they encounter a new host species^36, 37^. The maximum pairwise distance between cichlids in Lake Tanganyika is roughly equivalent to that of human and chimpanzee, with some species having pairwise distances of less than 0.03%, which are among the lowest values observed in all vertebrate groups^22^. It is noteworthy that the genetic distance between humans and chimpanzees is often sufficient for successful cross-species virus transmission^38^. This suggests that the close genetic relationships among cichlids, in line with their rapid speciation, should provide viruses access to a broad spectrum of cichlid hosts within Lake Tanganyika. A case in point is the presence of African cichlid picornavirus in five species that are members of four tribes: the Lamprologini, Ectodini, Perissodini, and Tropheini (Figure 4).

It is also likely that African cichlids will have similar immune properties, increasing virus susceptibility, and are frequently exposed to viruses. For example, a 125 m^2^ area in Lake Tanganyika can harbour up to 39 different cichlid species^39^. In contrast, reef fishes—that are characterised by multiple families with greater genetic diversity—show minimal virus cross-species transmission despite ample exposure among species^10^. Indeed, a recent study found that only four of the 38 viruses identified in a spatially restricted—100 m^2^—coral reef fish community, comprising 60 fish species (16 families), were associated with host-switching, and only on relatively deep evolutionary timescales^10^. Moreover, in this reef system fish are consistently found in groups of up to 25 species in a 3.5 m^2^ area, such that they represent the largest species densities among all extant vertebrate species^40^. Despite such substantially high rates of contact and exposure, the lack of host-switching in the reef ecosystem implies that there are strong host barriers to infection, particularly given the large degree of genetic divergence among reef fish families. Indeed, the cross-species transmission events observed were only identified in members of the same fish family, the Gobiidae. This further supports the notion that close genetic relatedness is a key determinant of rapid virus host-switching in African cichlids.

We assembled a time-calibrated phylogeny of all representative ray-finned fish hepaciviruses. It was notable that the rate of hepacivirus evolution was not constant throughout the cichlid radiation, but increased rapidly during the last 2-3 Ma, coinciding with a period of rapid diversification, packing of niche space and micro-niche partitioning in Tanganyikan cichlids^20^. This strongly suggests that as cichlid communities became denser and more diverse, viruses exploited a larger number of closely related host species for infection through cross-species transmission.

As well as hepaciviruses, our analysis identified 34 hepadnaviruses, of which 97% were closely related, indicative of recent cross-species transmission (Figure 4). In particular, a novel group of metahepadnaviruses fell sister to *Astatotilapia* metahepadnavirus that was recently discovered in *Astatotilapia* species from Lake Masoko. *Astatotilapia* belongs to the Haplochromini, which is the most species-rich clade of cichlids and widespread in Africa, where it diversified in many rivers and lakes including Lake Malawi^21^. This suggests that cichlid metahepadnaviruses are widely distributed in East African freshwater systems and have likely diversified alongside cichlids during multiple radiation events. A recent molecular clock analysis of the *Hepadnaviridae* estimated that the genus *Metahepadnavirus*, characterised exclusively by ray-finned fish hosts, split from orthohepadnaviruses, a group of mammalian viruses (e.g., hepatitis B virus), approximately 240 Ma, with both *Astatotilapia* and Mexican tetra (*Astyanax mexicanus*) metahepadnaviruses diverging ∼80-100 Ma^30^. However, the Haplochromini formed in Lake Tanganyika ∼5.5 Ma, and *A. mexicanus*, both surface dwelling and cave populations, originated very recently during the late Pleistocene, approximately 20 thousand years ago^41^. This strongly suggests that metahepadnaviruses have likely diverged much more recently than previously estimated^30^.

We identified piscichuviruses with high abundances in the brain tissue of three cichlid species. Currently, the genus *Piscichuvirus* (*Chuviridae*) is described by viruses associated with ectothermic vertebrates such as snakes, turtles, and ray-finned fish, as well as decapods^42^. Similarly, these viruses were recently detected in the brain and spinal cord of boa constrictors and three aquatic turtle species, causing lymphocytic meningoencephalomyelitis^43, 44^. Despite the divergence between fish and reptile piscichuviruses (∼48% similarity), our findings show that this emerging group is localised to the central nervous system and have likely maintained this tropism over millions of years.

Given their exposure to the external environment, it was not surprising that across all virus groups, the highest number of viral transcripts were identified in the gills and the LPJ, which together share a developmental origin and exhibit similar patterns of gene expression^25^. Conversely, we detected a very low number of viral transcripts in the gonads. This suggests that the viruses in this ecosystem might utilise cell receptors that aren’t naturally expressed in cichlid reproductive organs. It is also possible that the gonads might contain a rich source of antimicrobial peptides, as observed in the seminal fluid proteome of sparrows^45^. Indeed, it was recently shown that the rate of transcriptome evolution and the number of both protein-coding genes and long noncoding (lnc) RNAs are higher in the gonads compared to all other cichlid organs examined in this study^25^. Transcriptome evolution in the gonads should therefore experience strong selective pressures given their function in sperm competition and mate choice, and perhaps might select for antiviral mechanisms that protect the gametes from infection^46^.

While our analysis predicted multiple host-switching events and potential tissue tropisms, these are necessarily based on sequence analysis alone, such that the precise mechanisms of infection and the mode of transmission are not covered here and require further investigation. This is of particular importance given the geographical proximity and evolutionary relatedness to one of the most farmed and consumed fish species globally, the Nile tilapia, that often experiences devastating impacts from emerging viral infections^47^. Indeed, because the viruses identified are novel, their receptors are unknown. Moreover, the transcriptome data screened were gene expression profiles generated for host evolutionary analyses only^25^ and therefore not optimised for virus discovery. Nevertheless, we discovered a large diversity of viruses in 72% of the cichlid species examined.

Taken together, we show that the African cichlid radiation has had a profound impact on the extent, pattern, and evolution of virus diversity in Lake Tanganyika. While virus-host co-divergence is a common macroevolutionary pattern that forms the “backbone” of many family-level virus phylogenies, in the context of a host adaptive radiation virus diversification is accelerated by frequent cross-species transmission, especially during a time of rapid host diversification and increased niche packing in Lake Tanganyika^19, 20^. Hence, the rich viral diversity shared by the cichlids is likely the product of close genetic relatedness in combination with frequent interactions in Lake Tanganyika.

## Methods

### Assembly of cichlid viromes

Brain, liver, gill, lower pharyngeal jaw, ovary and testis transcriptomes from 74 Tanganyikan cichlid species (n = 445 individuals, 2242 RNA sequencing libraries) were obtained from a previous study^25^ (deposited at the NCBI Sequence Read Archive under BioProject PRJNA552202). Raw reads were quality trimmed using Trimmomatic (v.0.38)^48^ with parameters SLIDINGWINDOW:4:5, LEADING:5, TRAILING:5, and MINLEN:25 and then *de novo* assembled into contigs using MEGAHIT (v.1.2.9)^49^, with default parameter settings. Assembled contigs were compared against the protein version of the Reference Viral Database (RVDB) (August 2022), NCBI non-redundant protein (nr) and nucleotide (nt) databases (August 2022) using DIAMOND (BLASTX) (v.2.0.9) and BLASTn^50, 51^. To enable the identification of divergent viral sequences, we used an e-value search threshold of 1 × 10^−5^ as in previous studies^2, 10–13^. Contigs with top matches to the kingdom “Viruses” (NCBI taxid: 10239) were predicted as open reading frames (ORFs) using Geneious Prime (v.2022.0) (www.geneious.com)^52^. All putative viral ORFs were translated into amino acid sequences and used as a query to perform a second search (DIAMOND BLASTP) against the NCBI nr database to remove false positives. ORFs with top matches to fish or bacterial genes were deemed as probable false positives and removed from further analysis. Viral contig contamination and completion was determined using CheckV^53^. Contig abundances were calculated using RNA-Seq by Expectation Maximization (RSEM) (v.1.3.0) and coverage was assessed by mapping using Bowtie2 (v.2.3.3.1)^54, 55^. Virus abundance plots were generated using the phyloseq (v.1.42) and ggplot2 (v.3.3.6) packages in R (v.4.2.2)^56, 57^.

### Identification and removal of endogenous viral elements

To determine whether our putative viral contigs were expressed endogenous viral elements (EVEs) rather than exogenous viruses, we screened for: (i) disrupted ORFs and (ii) flanking host regions using CheckV and BLASTn^10, 53^. Contigs that contained intact ORFs with no flanking host genes were compared to all available cichlid genomes using TBLASTN with an e-value search threshold of 1 × 10^−20^. Sequences identified as EVEs were removed from further analysis.

### Taxonomic assignment of cichlid viruses

To taxonomically assign our newly discovered viruses, we aligned their predicted amino acid sequences (complete or partial genomes) with the complete genomes of related viruses available on NCBI/GenBank using MAFFT (v.7.450) (E-INS-i algorithm)^58^. To infer whether our viruses were novel species, we used levels of sequence similarity (e.g., amino acid p-distances) and phylogenetic grouping (see below) as specified by the International Committee of Viral Taxonomy (ICTV) (https://talk.ictvonline.org) for each viral genus/family. We also used these criteria to determine whether a virus was likely vertebrate-specific (i.e., infecting cichlid fishes) or associated with fish diet, microbiome or environment (i.e., “non-vertebrate”), particularly as viruses from these different major host groups are usually phylogenetically distinct^10, 11, 14^. Viruses identified as likely of non-vertebrate origin were excluded from further analysis.

### Annotation of cichlid viral genomes

Viral genomes were annotated with the ‘Live Annotate and Predict’ tool implemented in Geneious using related sequences from NCBI/GenBank, with a similarity threshold of 20%. To annotate specific protein domains, we used InterProScan with the TIGRFAMs (v.15.0), SFLD (v.4.0), PANTHER (v.15.0), SuperFamily (v.1.75), PROSITE (v.2022_01), CDD (v.3.18), Pfam (v.34.0), SMART (v.7.1), PRINTS (v.42.0), and CATH-Gene3D databases (v.4.3.0)^59^. To annotate divergent regions, we employed Phyre2 which uses advanced homology detection methods for protein structure prediction^60^. We used a confidence level of ≥95%.

### Phylogenetic analysis

To infer the evolutionary histories of cichlid viruses, we first used MAFFT (E-INS-i algorithm) to align the sequences determined here with representative background sequences for each viral family/subfamily/genus. In the case of RNA viruses, we utilised amino acid sequences of the most conserved RNA-dependent RNA polymerase (RdRp), while the DNA polymerase and capsid proteins were used for DNA viruses. Background sequences were selected from the ICTV and downloaded from NCBI/GenBank. Amino acid sequence alignments were trimmed using TrimAl (v.1.2) to remove ambiguously aligned regions with a gap threshold of 0.9 and a variable conserve value^61^. For each data set the best-fit model of amino acid substitution was estimated with the ‘ModelFinder Plus’ (-m MFP) flag in IQ-TREE^62, 63^. We used a maximum likelihood approach to estimate phylogenetic trees using IQ-TREE (version), with 1000 bootstrap replicates used to assess nodal support. All trees were annotated using FigTree (v.1.4.4) (http://tree.bio.ed.ac.uk/software%20/figtree/).

### Co-phylogenetic analysis of cross-species virus transmission and co-divergence

To visualise the relative frequency of cross-species virus transmission and virus-host codivergence within each viral group, we determined the cophylogenetic relationship between viruses and their hosts. A complete phylogeny based on genome wide data of Tanganyikan cichlids was downloaded^20^ and pruned to those taxa in which a virus was identified using the R packages phytools (v.1.0-3)^64^ and ape (v.5.6-2)^65^. A cladogram of the non-cichlid hosts was generated using the phyloT software, a phylogenetic tree generator based on NCBI taxonomy (http://phylot.biobyte.de/). The pruned cichlid and non-cichlid cladograms were then concatenated to form a host cladogram for each viral group. Tanglegrams that illustrate the link between host and virus trees were created using phytools and ape. The virus phylogenies used in the cophylogenies were constructed as described above. The relative frequencies of cross-species virus transmission versus virus-host codivergence were quantified using eMPRess^31^, which uses a maximum parsimony approach to determine the optimal “map” of the virus phylogeny onto the host phylogeny. The cost of duplication, host-jumping, and extinction event types were set to 1.0, whereas the cost of virus-host codivergence was set to zero since it is regarded as the null event. The number of generations and the population size for both virus and host were set to 100.

### Time calibration and diversification rates of cichlid hepaciviruses

We used BEAST2 (v.2.7.3)^66^ to estimate a time-calibrated phylogeny of representative ray-finned fish (Actinopterygii) hepaciviruses, utilising the NS5 amino acid sequence alignment (see above). To achieve this, we employed a Yule speciation model with a single, normally distributed calibration point at the root node of the cichlid hepacivirus clade based on prior age estimates of the formation of Lake Tanganyika (9-12 Ma) as well as the age of tMRCA of the cichlid radiation that was recently estimated to have diverged around 9.7 Ma^20, 23^. Our calibration was therefore set with a mean of 10.5 and standard deviation of 0.6 (interval 9-12 Ma). The best-fit evolutionary model was selected using BEAUti (v.2.7.3)^67^. We tested a combination of JTT, WAG, and LG amino acid substitution models with strict and relaxed clocks (log-normal or gamma). Based on tree topology, Effective Sample Size (ESS) and marginal likelihoods, a strict, gamma clock with the LG substitution model, a gamma distribution of among-site rate variation and a proportion invariant of 0.077 were identified as the most appropriate model. We ran the model using a Markov chain Monte Carlo (MCMC) chain with 50 million iterations, sampling every 10000, with convergence and ESS assessed using Tracer (v.1.7.2)^68^. The maximum clade credibility (MCC) tree was constructed using TreeAnnotator (v.2.7.3) after a 10% burn-in protocol (https://www.beast2.org/treeannotator/). Finally, we computed the gamma statistic to test the null hypothesis that the rate of hepacivirus diversification was constant over the cichlid radiation using the ape package in _R34,65._

### Phylogenetic and ecological diversity analysis

To determine whether virus transmission across Lake Tanganyika cichlids is random with respect to phylogenetic position and the ecology of the species, we evaluated the ecological and phylogenetic diversity of the infected and uninfected host species compared to a random subset. Phylogenetic diversity was calculated as the mean pairwise phylogenetic distances, which were extracted from the host tree using the cophenetic function in the R package ape (v.5.6-2)^65^. Ecological diversity was calculated as the mean pairwise Euclidean distance of available carbon (δ^13^C) and nitrogen (δ^15^N) stable-isotope measurements of the Tanganyika cichlid species^20^. We then assessed if the diversity of the infected and uninfected set of species (observed) is smaller than the diversity of a random subsample of the radiation (random) by randomly subsampling the total number of infected and uninfected species from the entire radiation 1000 times. If the observed diversity is lower (random-observed > 0) in more that 95% of the random samples (one-tailed test), we considered the difference significant (transmission and infectivity is assortative). This resampling approach was applied once across all viruses together, and once separately for each virus group that was present in at least five species in our data set.

## Supplementary Information

**Supplementary Table 1**. Description of the vertebrate-associated viruses identified in this study.

**Supplementary Table 2**. NCBI/GenBank accession numbers of the sequences used in the viral phylogenies.

**Supplementary Figure 1**. Time-calibrated phylogeny of the cichlid (host) adaptive radiation in Lake Tanganyika with host labels. The phylogeny was obtained from a previous publication^20^ and pruned to the species examined in this study.

**Supplementary Figure 2**. Phylogenetic (A) and ecological diversity (B) of infected (red) and uninfected (blue) species compared to the overall (grey) diversity of cichlids in Lake Tanganyika. C) Violin plots of the difference between the diversity (mean pairwise phylogenetic and ecological distances) of random sets of species (1000 subsamples of the radiation) minus the diversity of the observed set of infected and uninfected species, respectively.

**Supplementary Figure 3**. Phylogenies of the *Paramyxoviridae*, *Nidovirales*, *Hamaparvovirinae* (*Parvoviridae*), and *Arenaviridae*. Trees were estimated using amino acid sequences of the RdRp gene for RNA viruses and NS1 gene for the parvoviruses. Trees were midpoint rooted for clarity only. Circles on branch tips represent individual viruses identified in this study and coloured according to tribe. The scale bar represents amino acid substitutions per site.

## Acknowledgements

This work was funded by an Australian Research Council Discovery Project (DP200102351) and a National Health and Medical Research Council Investigator grant (GNT2017197) to ECH, and the Swiss National Science Foundation to FR (206869) and WS (208002). We acknowledge the University of Sydney for providing the high-performance computer cluster “Artemis”, that was used for this study.

## Author contributions

VAC, FR, WS and ECH conceptualised the study. VAC, FR, and JCOM performed the analyses. VAC wrote and prepared the original draft. VAC, JCOM, ECH, EH, FR, and WS edited and revised the manuscript. ECH funded the project. EH and ECH supervised the project.

## Data availability

The RNA sequencing reads used in this study were publicly available data on the NCBI SRA under BioProject PRJNA552202. All virus sequences have been deposited in NCBI/GenBank under the accessions XXXX-XXXX.

