## Supplementary figures and images for "Host adaptive radiation is associated with rapid virus diversification and cross-species transmission in African cichlid fishes"

### Supplementary Figures

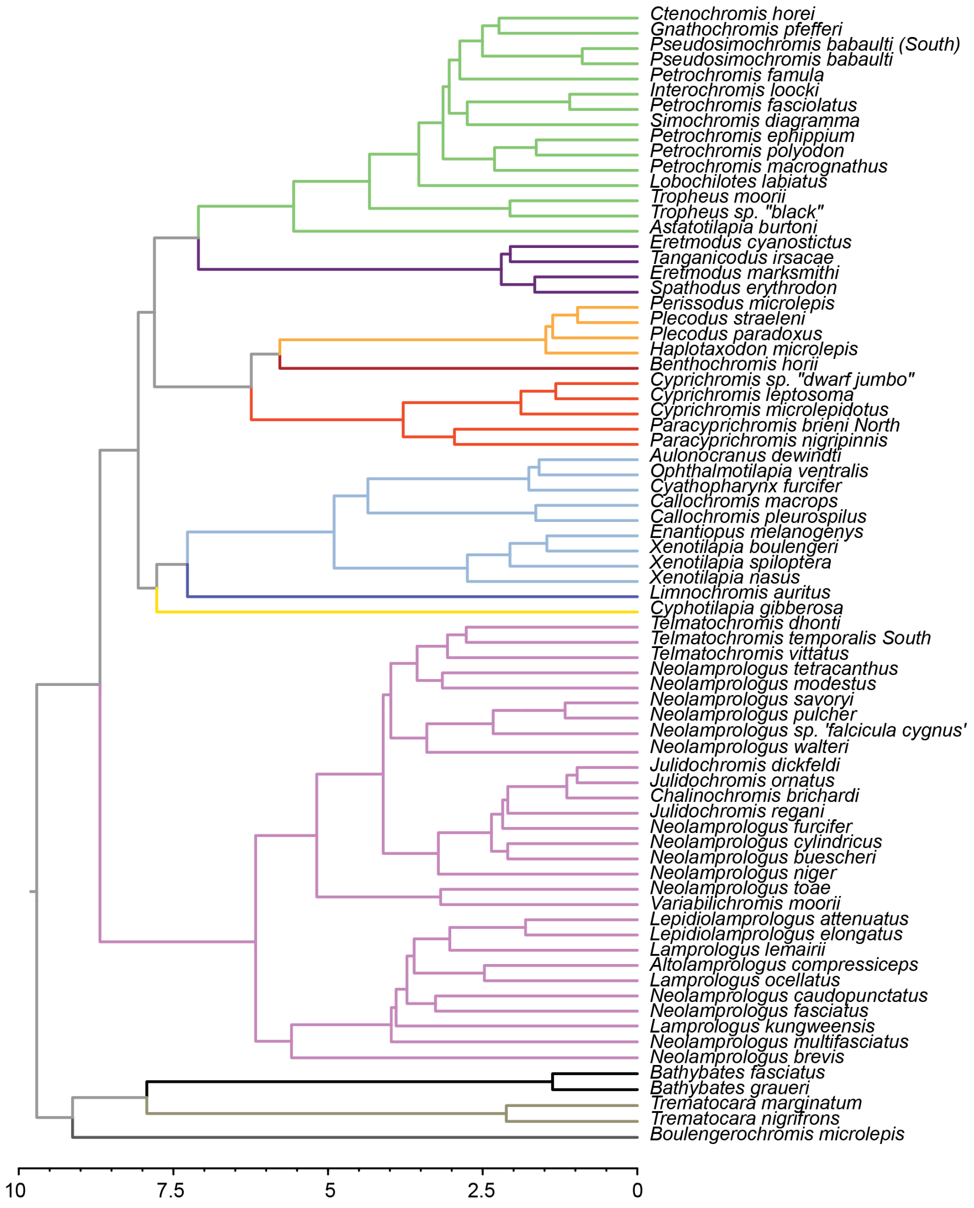


**Supplementary Figure 1**

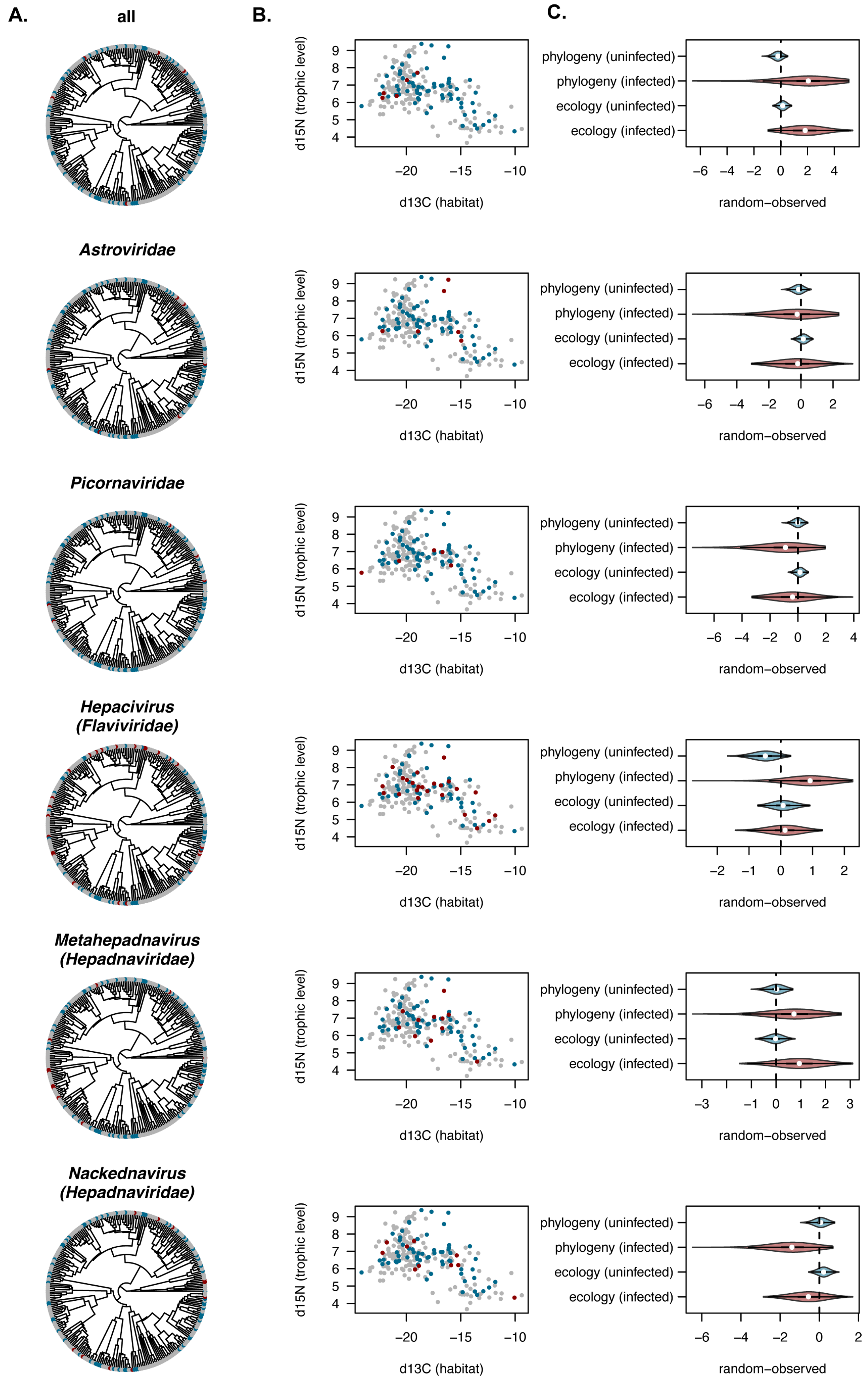


**Supplementary Figure 2**

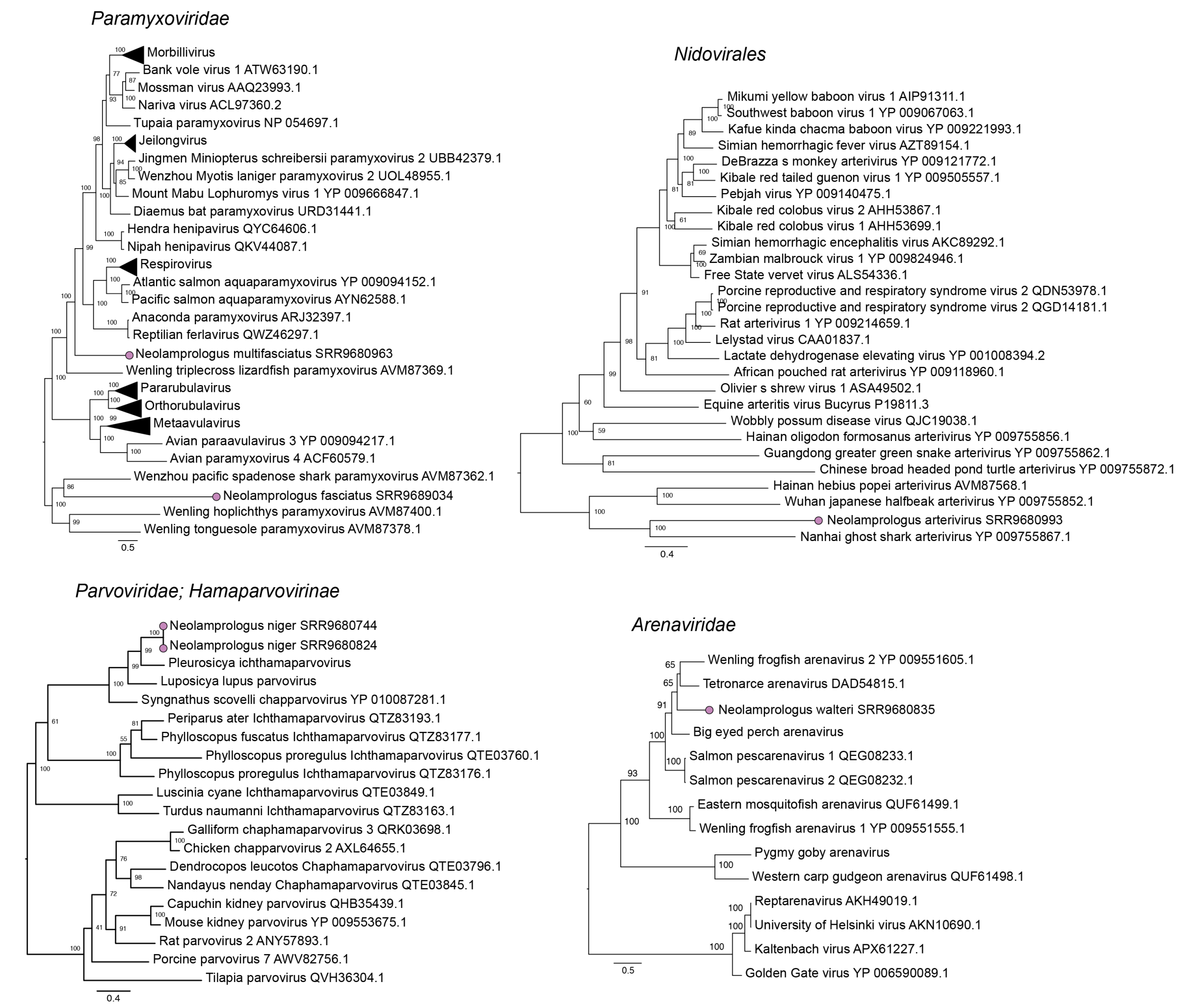


#### Supplementary Figure 3
